# HIV-1 and *Chlamydia trachomatis* restrict their respective growth but promote their survival in co-infected human macrophages

**DOI:** 10.1101/2025.08.19.671145

**Authors:** Mariano Alonso Bivou, Floriane Herit, Thibault Leray, Maria-Teresa Damiani, Florence Niedergang

## Abstract

HIV-1 and Chlamydia trachomatis (CT) are two significant sexually transmitted pathogens that frequently co-infect individuals. However, the mechanisms by which these two obligate intracellular pathogens interact at the cellular level remain elusive, particularly in tissue macrophages, where persistent infections can occur. In this study, we demonstrate that CT generates inclusions in macrophages of murine and human origins. We also show that both HIV-1 and CT reciprocally restrict the growth and replication of each other within co-infected human macrophages, irrespective of whether the viral or bacterial infection is established first. Notably, the co-infection resulted in improved survival of the macrophage hosts, as the inflammatory cell death pathways induced by CT were prevented by the virus. Collectively, these findings demonstrate that HIV-1 and CT collaborate to persist in human macrophages.

**IMPORTANCE:** While HIV-1 and *Chlamydia trachomatis* (CT) infections are associated at the epidemiological level, very little is known at the cellular and molecular level on co-infections by these two intracellular pathogens. The significance of our research is in dissecting the impact of one pathogen on replication and production of infectious progeny by the other. For this, we studied human macrophages, which are targeted by both HIV-1 and CT and could play an important role in their intracellular persistence. This work reveals the close interplay between these two pathogens that benefit from each other to survive in human macrophages. Consequently, we emphasize the need to address these cells as a unique target during the co-infections.

## INTRODUCTION

*Chlamydia trachomatis* (CT) is the most frequent bacterial cause of sexually transmitted infections (STIs) with 128.5 million new infections detected annually (1). The global prevalence of chlamydial infection has been estimated at 3.8 % in women and at 2.7 % in men, ranging from 1.5 % in developed countries to 6 % in developing countries (2).

The human immunodeficiency type 1 virus (HIV-1) still infects 39 million people worldwide and the number of new cases each year is estimated at 1.3 million adults. Co-infections between these two sexually transmitted pathogens are common. It is estimated that gonorrhea and chlamydia infections account for 3 % to 20 % of HIV transmission and 2 % to 15 % of HIV acquisition (3, 4). Both HIV-1 and CT are highly adapted sexually transmitted human pathogens, can be transmitted to newborns, and have obligate intracellular development. However, little is known about the molecular and cellular mechanisms underlying the high frequency of the epidemiologically observed association between CT and HIV infection.

Both HIV-1 and CT can enter a persistent state and survive within infected cells for years; therefore, their infections can evolve to a chronic persistent course associated with chronic low-grade inflammation (5–7). CT persistence is defined by the temporary interruption of its developmental cycle to lead to a viable but non-infectious, non-replicative state. Persistence may be triggered by various stimuli, such as treatment with penicillin or INFγ, deprivation of tryptophan or sphingolipids or coinfection with herpes virus (6, 8). Upon removal of the stressor, CT re-enters its normal developmental cycle. Moreover, CT can manipulate intracellular signaling pathways within the infected cell to restrict the activation of the host immune response. Chlamydia was shown to activate inflammasomes to induce processing and release of the NF-κB-dependent IL-1β and IL-18, which leads to an inflammatory form of cell death termed pyroptosis (9, 10).

In addition, both pathogens affect immune cells functions (11–14). Indeed, numerous reports indicate that innate immune cells, such as macrophages, partially lose their functions because of HIV-1 infection (15). Our work has further revealed that several viral factors are implicated in the inhibition of the phagocytic and activation properties of these cells (16–18). Of note, the role of urethral macrophages as a reservoir of HIV-1 has been highlighted as they have been found even in patients under combined anti-retroviral therapy (cART) (19). The role of macrophages in chlamydial infections has long been overlooked. However, with the emergence of recent reports describing chlamydial antigens in extragenital tissues (20, 21), the outbreak of invasive lymphogranuloma venereum (LGV) infections since the beginning of this century (22) and the antibiotic resistance of CT when infecting macrophages (23), this immune cell has been repositioned as a key contributor in the transport and dissemination of viable bacteria in the body (24–26). Recent reports indicate that CT is capable of infecting macrophages in human patients (25, 27) and the presence of aberrant bacterial forms that account for persistent infections (20, 28, 29).

At the genitourinary tract level, CT infections induce inflammation and favor the development of co-infections. Indeed, alterations in mucosal barrier continuity secondary to chlamydial infection promote HIV-1 infection and vice versa (30–33). Furthermore, the release of proinflammatory mediators, by recruiting immune cells, promotes the establishment of infections.

In this work, we observed that primary human macrophages may harbor HIV-1 and CT infections simultaneously. We documented that co-infection had a mutual delaying effect on the replicative cycle of both pathogens, while altering the host signaling pathways and thus, the inflammatory and cell death response of the macrophages in favor of creating a niche where both pathogens can persist.

## RESULTS

### Macrophages support *Chlamydia trachomatis* lifecycle and productive infection

CT is a highly adapted human pathogen that primarily infects epithelial cells. Recently, infections of myeloid cells have been described (22), thus, we assessed the ability of CT to grow within macrophages. We comparatively infected a murine macrophage cell line, the RAW264.7 cells, and primary human monocyte-derived macrophages (hMDM). Not only did CT successfully invade both types of macrophages, but it also developed in both cell models, providing further evidence of the bacterial ability to survive within professional phagocytes, human or murine (Fig 1A). As we aimed to study the CT/HIV coinfection, we further described the chlamydial lifecycle within human macrophages, the natural tissue niche where HIV-1 persists (15). We infected hMDM for 24 h, 48 h, and 72 h with GFP-CT and analyzed infected cells by fluorescence microscopy. CT529, a chlamydial protein that decorates the membrane of CT-containing vesicles (inclusions), showed a typical pattern of expression which confirms inclusion integrity within the phagocyte (Fig 1B). Individual CT invade the host cell in a plasma membrane-derived compartments that travel towards the perinuclear region where they fuse homotypically to form a single compartment within the first 24 h post-infection (pi) in epithelial cells (35). In hMDM, we observed that more than 70 % of infected cells displayed two or more inclusions per cell at 24 h pi, while the cells bore a single inclusion only after 72 h, indicating delayed fusion dynamics as compared with the kinetics reported in epithelial cells (Fig 1C). However, individual inclusion area was assessed over time, showing that they can grow within macrophages (Fig 1D). Bacterial development comprises not only the growth of the inclusion, but also the differentiation of bacterial cells from a replicative un-infective reticulate body (RB) to a non-replicative infective elementary body (EB). Inclusion forming unit (IFU) was calculated in relation to the number of progenitor inclusions (INPUT) to compare the efficiency of RB to EB transition and replication occurring in other cell types. IFU quantification indicated that CT could replicate and differentiate within macrophages, accomplishing the entire lifecycle with comparable efficacy to that in epithelial cells (36) (Fig 1E).

**Figure 1:**
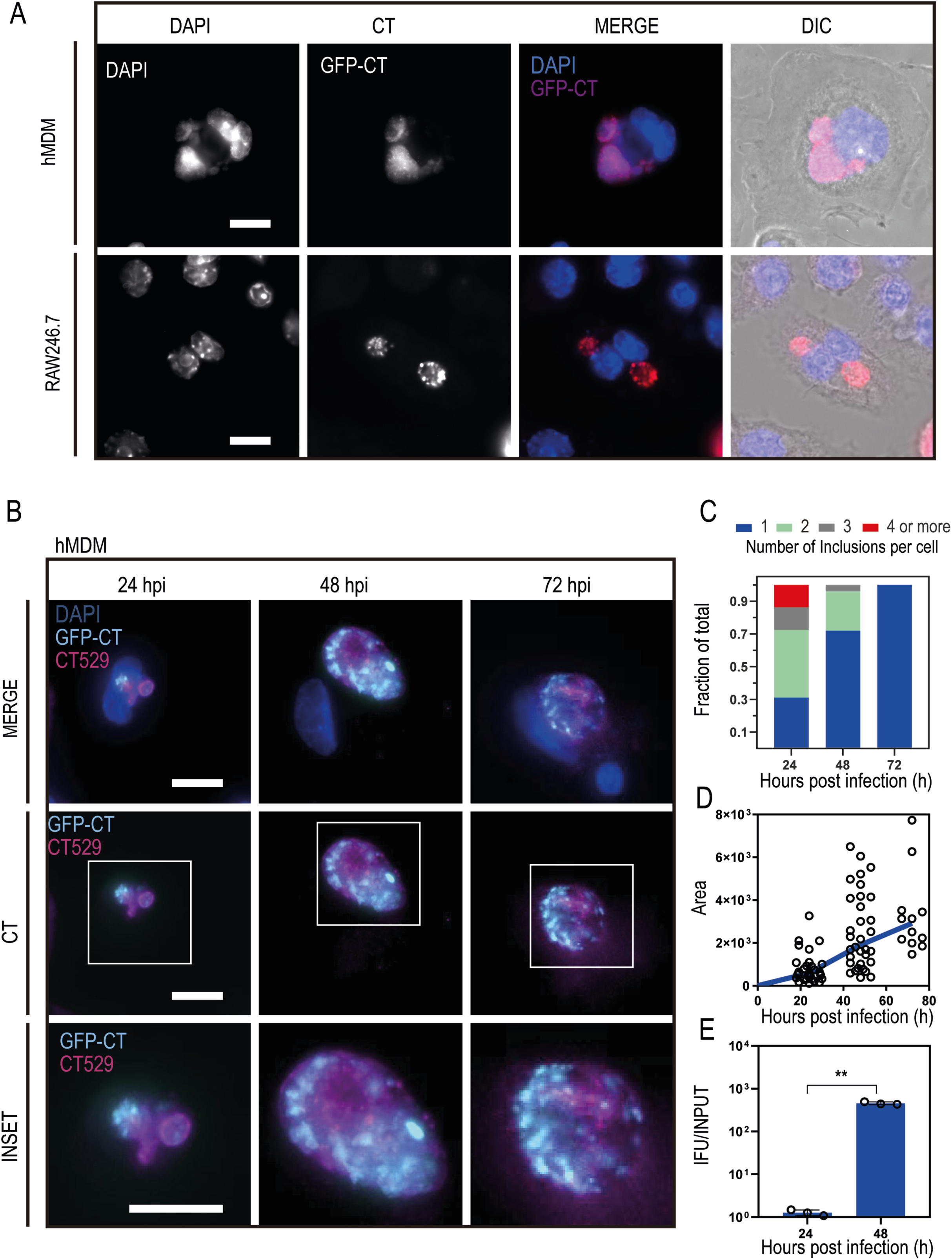
CT develops functional inclusions within human and murine macrophages. (A) Representative images of human (hMDM) and murine (RAW264.7) macrophages infected with GFP-expressing CT for 24 h. (B) Macrophages were infected for 24, 48 and 72 h. Chlamydial inclusions were detected with an anti-CT529. (A) and (B) DAPI was used to detect bacterial and eukaryotic DNA. (C) Quantification of the number of inclusions within individual macrophages after 24, 48 and 72 h of infection. (D) Area of individual inclusions at 24, 48 and 72 h pi. (E) Inclusion Forming Units (IFU) related to the INPUT at 24 and 48 h pi, representing three independent experiments. Data were analyzed using a Student’s t-test (** indicates p <0.01).

### HIV-1-superinfection of CT-infected macrophages restricts pathogen replication and reduces infective progeny

To study HIV-1 and CT co-infection, differentiated macrophages were infected with CT overnight (12 h) and then infected with HIV-1 for 24, 48 and 72 h (Fig 2A). A representative coinfected cell for 24 h is shown in Fig 2B. Inclusions can be observed with DAPI staining of their genetic material while HIV particles were immunolabeled with anti-capsid (CA)p24 antibodies. HIV and CT were always detected in two different pathogen-containing compartments, suggesting that heterotypic fusion does not occur at the observed time points. CT lifecycle progression was evaluated by IFU analysis at the three different time points in cells infected with CT, and then HIV-1- or mock-co-infected (Fig 2C). The analysis revealed that HIV-1 reduced the efficacy of CT transition from RB to EB only at 72 h pi, suggesting that HIV-1 has to establish a productive infection to affect the CT differentiation to EB in macrophages. In addition, the ability of HIV-1 to thrive within CT-infected macrophages was evaluated by the expression of the viral protein CAp24 in macrophage lysates that were CT- or Mock-infected (Fig 2D). Expression of CAp24 by ELISA was significantly reduced after 72 h when the macrophages were previously infected with CT as compared with the Mock-infected macrophages. A significant decrease in the secretion of viral CAp24 was detected even earlier, at 48 h pi, in the supernatant of infected macrophages (Fig 2E). Hence, we next evaluated the ability of the viral particles produced under both conditions to infect human cells using the TZM-bl assay (Fig 2F). The number of infectious viral particles was dramatically reduced in macrophages pre-infected with CT, as compared with mock-treated macrophages. Interestingly, immunofluorescence analysis of the expression of the viral CAp24 in CT- and CT-Mock-infected macrophages (Fig 2G) showed that CT-challenged macrophages were not as permissive for HIV replication as CT-Mock-infected cells, independently of inclusions development. Thus, both pathogens are able to infect the same cell and complete their lifecycle while reciprocally reducing their efficacy in the production of infectious progeny.

**Figure 2:**
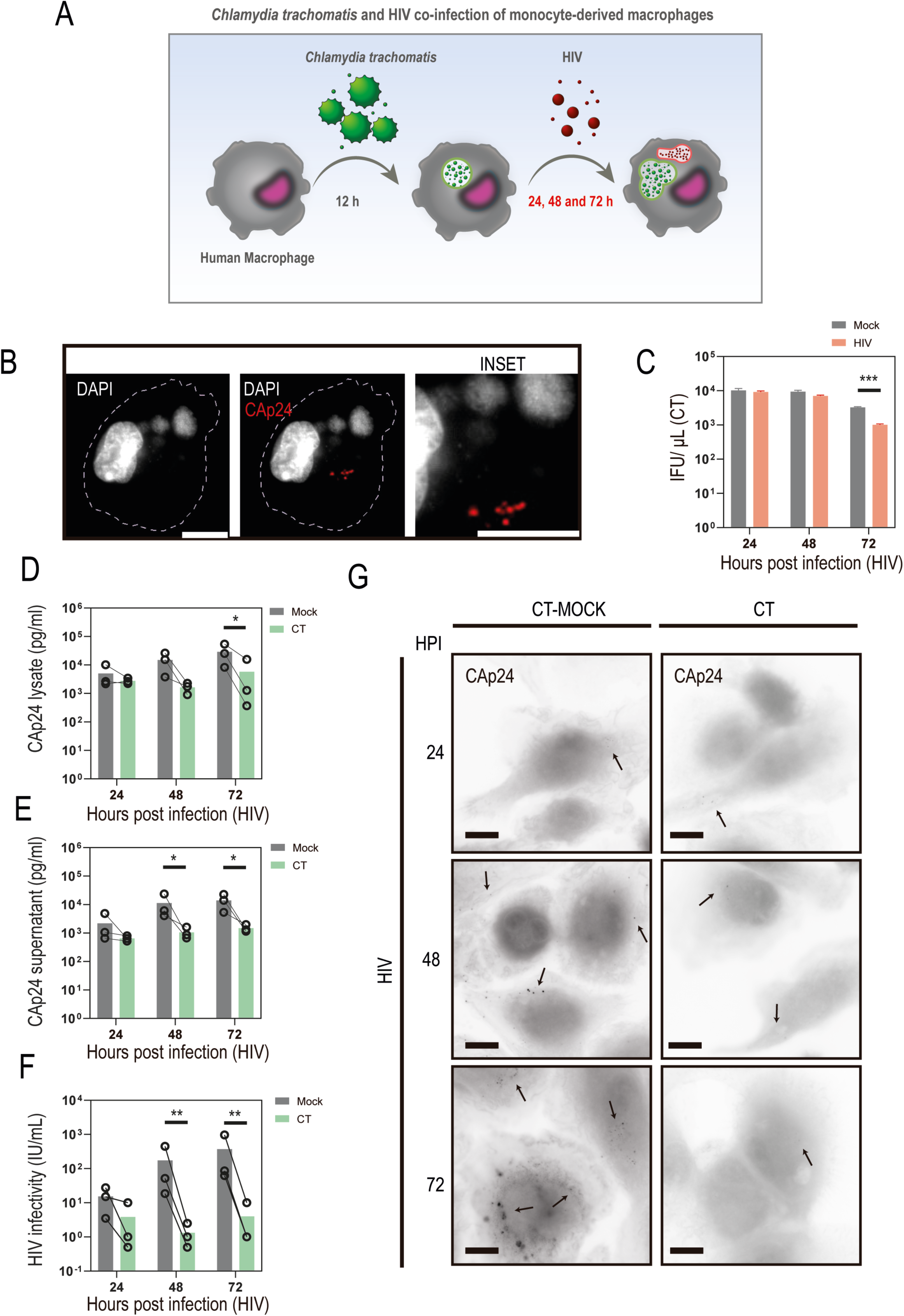
CT and HIV can reside within the same macrophage. A) Schematic representation of the experimental design. B) Representative fluorescence image of a co-infected human macrophage. HIV-1 CAp24 was immunodetected while human and CT DNA was stained with DAPI. C) Inclusion Forming Unit analysis of HIV and Mock CT-co-infected macrophages at different time points (24, 48 and 72 h pi). (D), (E), (F) and (G) Macrophages were CT or Mock infected before HIV-1 coinfection. Capsid CAp24 was quantified by ELISA in macrophage lysates (D) and in supernatants (E). (F) Infectivity of viral particles in supernatants was quantified by TZM-bl assay. (G) Immunofluorescence images of macrophages that were Mock or CT challenged but do not bear inclusions. Arrows mark HIV CAp24 positive structures within macrophages. (B) and (D) bars represent 10 μm. Graph in (C) shows mean ± SEM from n = 3 independent experiments and graphs in (D), (E) and (F) mean of n = 3 independent experiments. ∗p < 0.05, ∗∗p < 0.01 and ∗∗∗p < 0.001 for indicated comparisons from two-way ANOVA following adjustment for multiple comparisons.

### Chronic HIV-1 infection limits CT development in co-infected macrophages

HIV is able to replicate within macrophages without activating cell death programs, as opposed to the fate of HIV-1-infected lymphocytes (37, 38). It can therefore establish a chronic course of infection in these phagocytes. For this reason, we infected macrophages with HIV-1 or a Mock supernatant for 7 days before superinfecting them with CT for 24 or 48 h to further characterize the time-dependent aspect of their interaction (Fig 3A). From this point on, hours post-infection refer to the time between chlamydial challenge and samples collection. At 24 h pi, the chlamydial inclusions could be visualized in both the control and the HIV-1 infected conditions (Fig 3B). In this case, due to the higher viral load achieved during chronic infection, HIV-1 containing compartments were easily immunodetected at 24 h pi. Noteworthy, both pathogen-containing compartments remained unfused, but in close proximity. The size of the inclusions was measured by fluorescence microscopy, demonstrating that inclusions that developed within HIV-1 pre-infected macrophages reached a smaller area at 24 h pi (Fig 3C). At 48 h pi, the production of chlamydial infective particles was quantified by IFU analysis, confirming a reduction in bacterial development in cells already infected with HIV-1 (Fig 3D). Together, these results indicate a reduced CT development in macrophages pre-infected with HIV-1.

**Figure 3:**
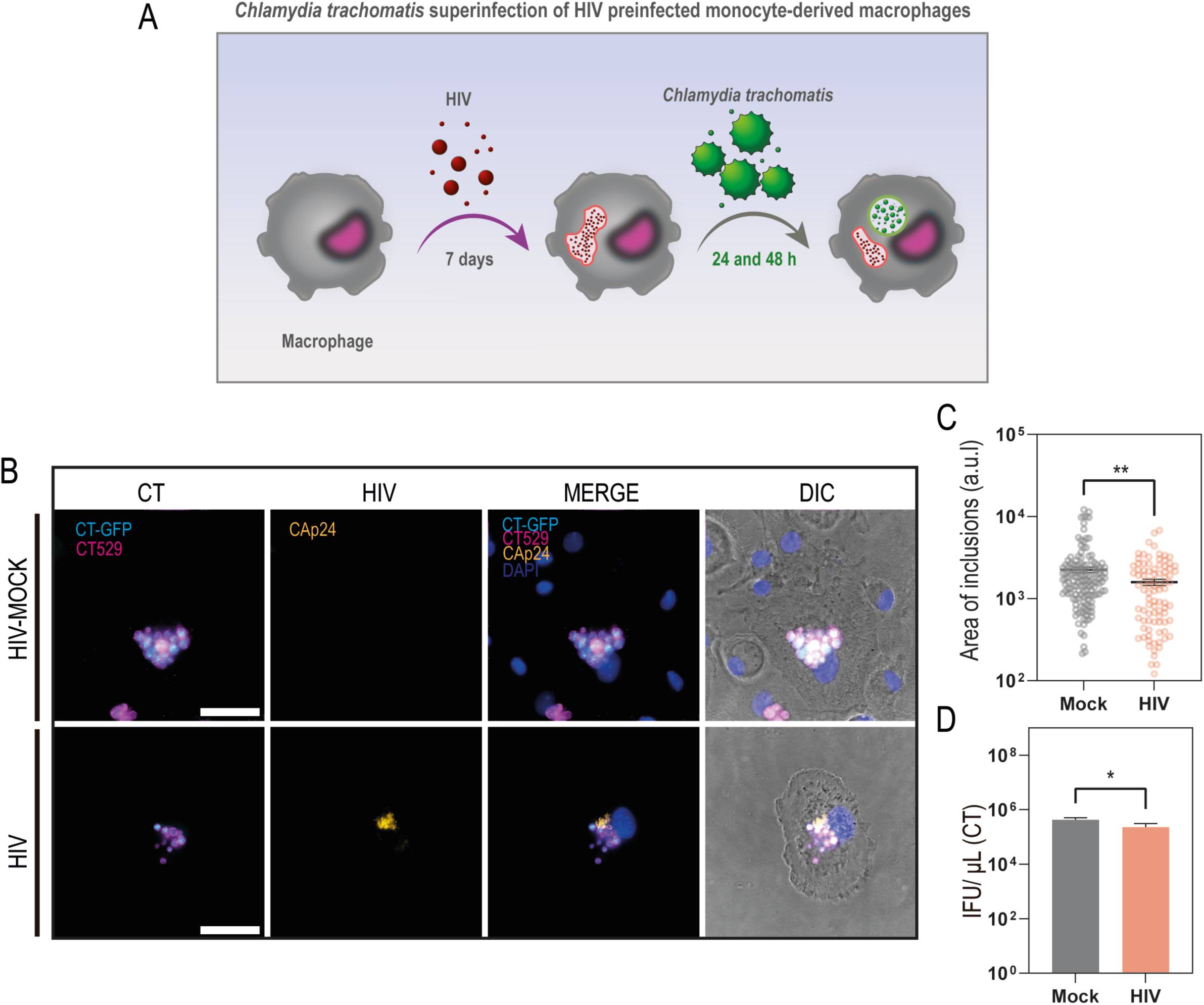
**Established HIV-1 infection of human macrophages limits CT development**. (A) Schematic representation of CT superinfection of HIV-1 infected macrophages. (B) Immunofluorescence representative micrographs of HIV and HIV-MOCK infected macrophages and then superinfected/infected with GFP-CT for 24 hours. (C) Area of individual inclusions superinfecting MOCK and HIV infected macrophages at 24 h pi. (D) IFU analysis of 48 h CT-infected macrophages that were previously MOCK or HIV infected. Graphs in (C) show mean ± SEM from representative of n = 3 independent experiments and in (D) n = 3 independent experiments. ∗p < 0.05 and ∗∗p < 0.01 for indicated comparisons from Student’s t test.

### CT superinfection of HIV-infected macrophages reduces viral burden

To study how chlamydial super-infection impacts on the development of an established HIV-1 infection, we used the same experimental design as before (Fig 3A). Fig 4A shows representative fluorescence images of HIV and CT-HIV infected cells (24 h pi). Macrophages infected for 8 days bear relatively large virus-containing compartments, as detected with an antibody against CAp24. After CT superinfection, these reservoirs were reduced in size as compared with Mock superinfection conditions (Fig 4A). The expression of CAp24 in co-infected cells was quantified by immunofluorescence confirming a decrease in viral burden in cells that support both pathogens (Fig 4B). By western blot, we also observed a decrease in the expression of CAp24 at 24 kDa and his precursor (Pr)55 at 55 kDa in the cell lysates of macrophages that were infected with CT for 24 and 48 h as compared with Mock-infected cells (Fig 4C). Additionally, quantification of intracellular CAp24 by ELISA confirmed that the CAp24 expression was significantly reduced at 48 h in CT-infected macrophages as compared to the corresponding Mock controls (Fig. 4D). In line with these results, the quantification of secreted CAp24 by ELISA showed that macrophages pre-infected with CT produced reduced levels of CAp24 in their supernatants at 48 h, as compared with CT-Mock-treated cells (Fig. 4E). Analysis of the infectivity of the viral particles by TZM-bl assay revealed that co-infected macrophages exhibited a strong reduction in their capacity to secrete infectious viral particles as early as 24 h pi, as compared with CT-Mock-infected macrophages (Fig. 4F). In conclusion, CT superinfection alters HIV-1 development in hMDM.

**Figure 4:**
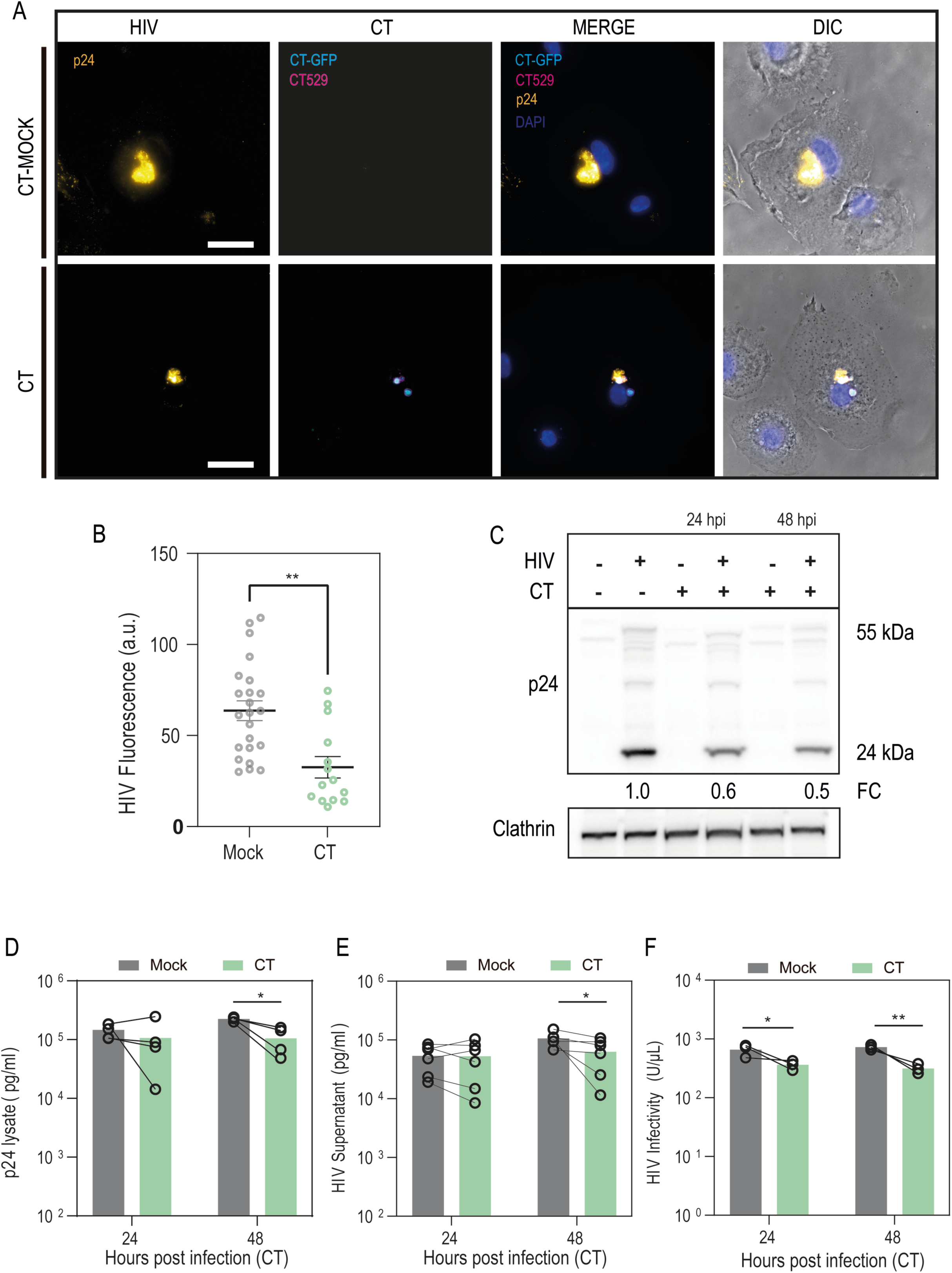
CT growth within HIV-1 pre-infected macrophages decreases viral load and production of infectious particles. (A) Representative immunofluorescence micrograph of HIV and HIV/CT-GFP coinfected macrophages. The viral protein CAp24 and the bacterial protein CT529 were immunodetected while DAPI stained DNA. hMDMs were infected with HIV for 7 days and then superinfected with CT for 24 (A) and (B), and 48 h (C), (D), (E) and (F). (B) Quantification of CAp24 expression by fluorescence microscopy. Fluorescence intensity of individual cells was analyzed with ImageJ after 24 h of CT infection. (C) HIV-1 p24 and p55 expression by western blot analysis after 24 and 48 h of CT infection. Clathrin was used a loading control. Quantification by ELISA of CAp24 in cell lysates (D) and in supernatants (E). (F) TZM-bl analysis of the infectivity of viral particles present in supernatants. Graph in (B) shows mean ± SEM from representative of n = 3 independent experiments and graphs in (D), (E) and (F) mean of n = 3 independent experiments. ∗p < 0.05 and ∗∗p < 0.01 for indicated comparisons from Student’s t test in (B) and from two-way ANOVA following adjustment for multiple comparisons in (D), (E) and (F).

### HIV-1 infection of macrophages prevents CT-induced inflammasome response and cell death

Because our previous results suggested that the production of infectious bacteria and viruses decreased when both pathogens infected the same cells, we sought to monitor the viability of the cells bearing these co-infections. While HIV-1 infection of macrophages is not reported to induce apoptotic signaling (39), Chlamydial antigens and metabolites detected by the host cells induce pyroptosis, an inflammatory cell death characterized by the processing and secretion of IL1β, which is dependent on the canonical and non-canonical activation of the inflammasome (9, 10, 40, 41). Thus, we analyzed the presence of the lactate dehydrogenase (LDH), an intracellular enzyme that is released in the extracellular medium upon necrosis or pyroptosis, in the supernatants of infected and coinfected macrophages as shown in Fig 3A. As expected, we observed an increase in LDH release over time upon CT infection, whereas HIV-1 infection alone did not induce any changes (Fig 5A). Remarkably, the LDH release induced by CT after 48 h was partially prevented in HIV-1 pre-infected cells. These results suggested that an established HIV-1 infection may counterbalance the cell death signaling induced by the CT infection.

**Figure 5.**
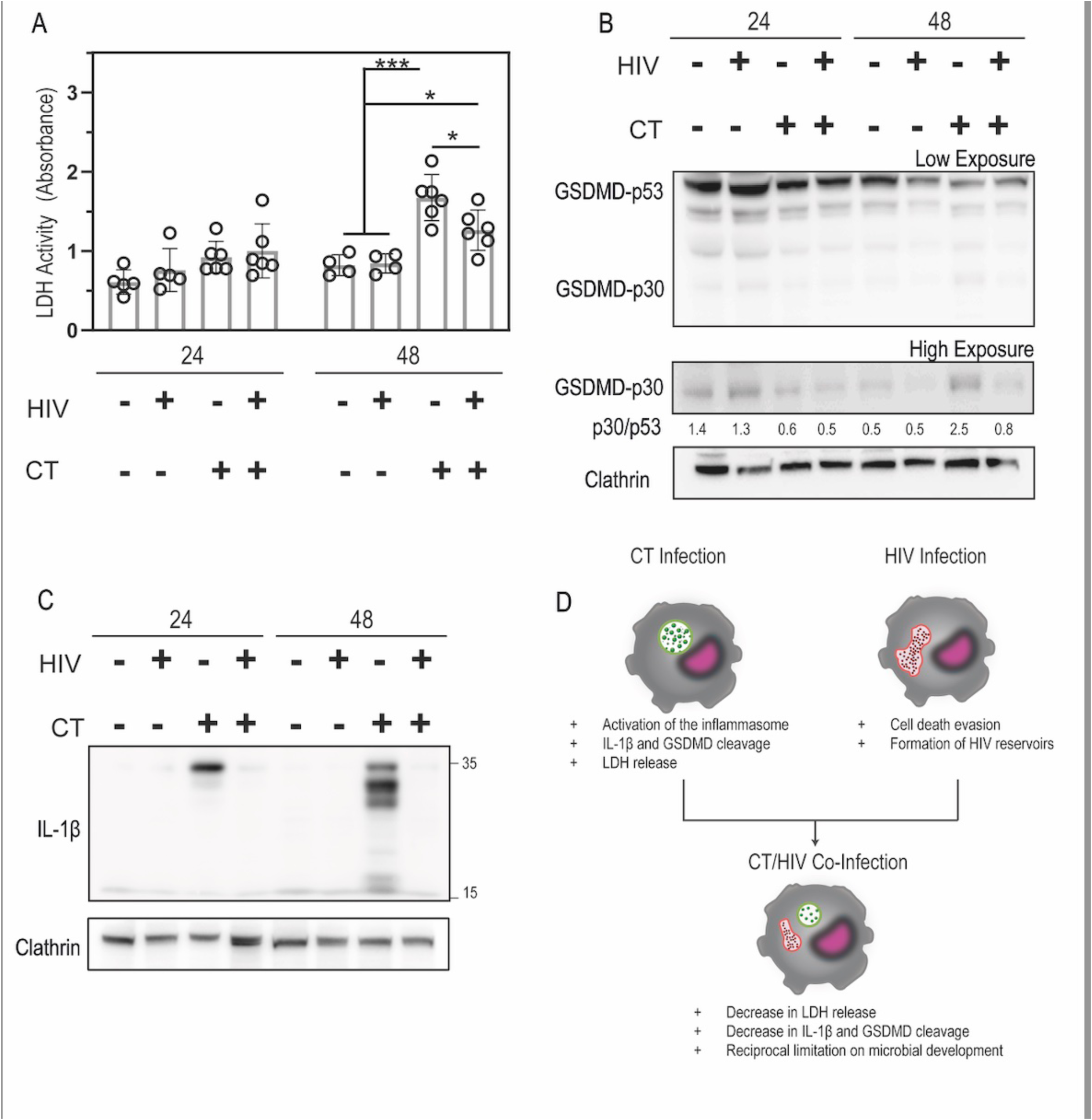
: Chronic HIV infection dampens inflammatory response to secondary CT infection. Macrophages were chronically infected with HIV for 7 days and later overinfected with CT for 24 and 48 h. (A) Cell viability was assessed by LDH release and activity in supernatants. (B) Expression of unprocessed and cleaved forms of GSDMD-p53 and GSDMD-p30, respectively. Clathrin was used as loading control. (C)Expression of unprocessed and cleaved forms of IL-1β of 31 kDa and 17 kDa, respectively. Clathrin was used as loading control. (D)Schematic summary of the proposed model Graph in (A) shows mean ± SEM from representative of n = 6 independent experiments. ∗p < 0.05, ∗∗p < 0.01 and ∗∗∗p < 0.001 for indicated comparisons from two-way ANOVA following adjustment for multiple comparisons.

To assess the activation of the inflammasome and pyroptosis pathway upon CT infection, we examined the processing of Gasdermin-D (GSDMD) (Fig 5B), which forms pores important for IL1β release (Fig 5C) and for the lytic process. A cleaved form of GSDMD was observed 48 h post-CT infection, consistent with the cell viability data (Fig 5A). Of note, the cleavage was much less pronounced in the HIV-1-preinfected cells. The expression of IL1β was induced after 24 h and 48 h of CT infection, and the cleavage of the protein was clearly visible after 48 h (Fig 5C). Strikingly, IL1β expression was absent or very weak in macrophages pre-infected with HIV-1 for 8 days prior to CT infection.

Together, these results indicate that an established HIV-1 infection of macrophages prior to CT infection prevents pyroptosis and cell death, which is beneficial for both intracellular pathogens.

## DISCUSSION

In this work we demonstrate that co-infection of human macrophages with HIV-1 and CT led to restriction in their growth and a better survival of the host cells, as the inflammatory cell death pathways induced by CT were prevented by the virus.

The role of macrophages in genital chlamydia infection in BALB/c mice was recently highlighted because these cells harbored the highest number of chlamydia DNA copies in the spleens of intravaginally infected mice (24). In another set of experiments, clodronate liposomes were used prior to CT infection to deplete the mice of their macrophages. Clodronate treatment not only decreased the overall bacterial load in the tissues, but also decreased the release of infective progeny at the genital level and the dissemination of the bacteria within the organism. Moreover, the use of TNF blockers inhibited the generation of inflammatory lesions in extra-genital tissues. In another study, they emphasize that even in the absence of direct CT infection of resident or recruited macrophages at the site of infection, these cells are the ones responsible for inducing an immunopathological response that sustains the pathophysiology of the infection (42). In addition, CT-infected macrophages were detected in human patients (25).

In this work, we present evidence that CT is able to generate inclusions in macrophages of both murine and human origins. Inclusions in human macrophages are able to grow in a pattern similar to that observed in epithelial cells. Furthermore, we show that CT is able to generate a number of inclusion-forming units per progenitor inclusion (INPUT) very similar to that observed in HeLa cells, demonstrating that CT is able to complete its growth cycle inside human macrophages with an efficiency that is similar to that observed in epithelial cells. This means that the bacterium successfully and globally modifies macrophage scavenging functions, including their vesicular trafficking, metabolic pathways and intracellular signaling events (24, 25, 43–46).

HIV-1 infection results in progressive immunodeficiency due to loss of CD4^+^ T lymphocytes and increased susceptibility to infections (47). ART restores CD4+ counts, prevents the development of Acquired Immuno-Deficiency Syndrome (AIDS) and has been effective in reducing mortality and morbidity in people with HIV (48, 49). Although treatment with ART eliminates the occurrence of opportunistic infections, the risk of developing bacterial and viral infections or cancer remains elevated in people living with HIV compared to the HIV-negative population (50–55). For some pathogens, the shared route of infection may be part of the reason for this increased risk, but residual immunodeficiency is still thought to be a central reason. The macrophage is currently considered to be one of the major reservoirs of HIV-1 *in vivo* and to be responsible for rebound and increased viremia after ART interruption (19, 56, 57). Therefore, an *in vitro* study of co-infection in a primary culture model of human macrophages is relevant. Using primary human macrophages, we show that co-infection is possible, and that both pathogens are able to replicate and produce infectious progeny within this cell type.

Upon infection with CT, macrophages are able to initiate inflammasome activation and subsequent pyroptotic cell death (9, 41). What was striking in our study was that CT superinfection in previously and chronically HIV-infected macrophages did not promote inflammasome activation, IL-1β induction and processing, and LDH release. In the case of CT and HIV co-infection, we observed limited growth of both pathogens regardless of the order of infection, but we still obtained infectious progeny of both CT and HIV. Interestingly, the outcome of the HIV-1 and CT co-infection differs from the infection with other bacteria like *E. coli* or *Salmonella* Typhimurium, which benefit from the impaired degradative capacities of HIV-1 infected macrophages and survive better in these cells (15, 16, 18). As an obligate intracellular bacterium, CT may need to compete with HIV-1 for resources, which could explain the mutual restriction in their growth during co-infections that we observed.

These results take on particular significance when one considers the manner in which Lymphogranuloma venereum, an infection caused by CT serovars L1-L3, has re-emerged since the turn of this century (58). This highly inflammatory infection had virtually disappeared from the Western world with the advent of antibiotics. However, in the last 20 years or so, and following the HIV pandemic, a large number of LGV cases with a very high association with HIV infection have been reported worldwide (22, 59–62). In particular, LGV in people living with HIV-1 display a less inflammatory clinical presentation with proctitis and proctocolitis (76), reinforcing a possible mutually beneficial relationship between HIV-1 and CT. Our findings show for the first time that both pathogens not only develop their intracellular niche but also generate infectious progeny within co-infected macrophages. In summary, HIV and CT display a restrictive growth yet cooperative survival in co-infected human macrophages, which appears as a valuable target to focus on in further studies on therapeutic strategies.

## MATERIALS AND METHODS

### hMDM differentiation and culture

Human monocytes were isolated from peripheral blood of healthy donors (Etablissement Français du Sang Ile-de-France, Site Trinité, Inserm agreement #18/EFS/030 ensuring that all donors gave a written informed consent and providing anonymized samples) by density gradient sedimentation on Ficoll (GE Healthcare), followed by adhesion to plastic at 37 °C for 2 h in the presence of adhesion medium (RPMI 1640 (Life Technologies) supplemented with 100 μg/ml streptomycin/penicillin and 2 mM L-glutamine (Invitrogen/Gibco). Then, the adhered cells were washed once with warm adhesion medium and differentiated in macrophage medium (RPMI 1640 supplemented with 10% FCS (Eurobio), 100 μg/ml streptomycin/penicillin, 2 mM L-glutamine) and 10 ng/mL recombinant human macrophage colony-stimulating factor (rhM-CSF; R&D systems) for 6-7 days.

### Infection and Propagation of CT

i) C. trachomatis Lymphogranuloma venereum, Type II was from ATCC (L2/434/Bu VR-902B) and the fluorescent strains p2TK2-SW2 IncDProm-RSGFP-IncDTerm (GFP-Ct) and p2TK2-SW2 IncDProm-mCherry-IncDTerm (mCherry-Ct) prepared by Hervé Agaisse and Isabelle Derré (Department of Microbial Pathogenesis, Yale University School of Medicine, New Haven, Connecticut, United States of America) were kindly given by Agathe Subtil (Institut Pasteur, Paris).

Infections were performed by adding a previously titrated suspension of purified EBs to the culture medium of the cells to be infected. The Multiplicity of Infection (MOI) is calculated as the number of infectious particles present in the suspension divided by the number of total cells susceptible to infection. It indicates the average number of infectious units per cell. After addition of bacteria, centrifugation was performed at 2000 × g, 10° C, 30 min. Thereafter, infected cells were washed with PBS and transferred to a culture incubator at 37 °C and 5 % CO_2_ for the indicated hours after infection (h pi).

For bacterial propagation, the HeLa cells were infected at a MOI of 2-5 and incubated for 48 h. Then, infected cells were lysed and EBs were purified on a density gradient as described previously (63). Purified EBs were suspended in 0.2 M sucrose, L-glutamine and phosphate buffer (SPG) (pH=7.2) and their content was titrated by quantification of inclusion forming units (IFUs) using confocal microscopy and/or flow cytometry. Stocks were stored in SPG buffer at -80°C.

### Production of HIV-1 particles

Proviral infectious clones of the macrophage-tropic virus isolate ADA (HIV-1_ADA_WT) have been described previously (18). Stocks of viral particles were obtained by transfection of HEK293T cells (Human Embryonic Kidney 293, ATCC®CRL-1573™, 2x10^6^) with 6 µg of the corresponding proviral DNA, using FuGENE® 6 Transfection Reagent as recommended by the manufacturer (Promega). Supernatants of the transfected cells were collected after 48 h, filtered, and stored at -80°C. Viral titers were assessed by infection of the indicator cells, HeLa TZM-bl (bearing the β-galactosidase gene under the control of HIV-1 LTR, National Institutes of Health-NIH reagent program), with serial dilutions of the stock, followed by a β-galactosidase coloration of the cells and counting of blue cells.

At 6-7 days of macrophage differentiation, HIV-1 (MOI 0.2) was added to human monocyte-derived macrophages (hMDM). Excess virus was removed after 2 days.

### Western Blot

Cells were lysed in lysis buffer (Tris HCL pH 7.5 20mM, NaCl 150 mM, NP40 0.5%, Protease Inhibitor, NaF 50mM, Sodium Orthovanadate 1mM) and scraped with a bent tip to collect the lysates. The lysates were collected in microtubes and centrifuged at 10000 g for 15 min at 4°C. Then, the supernatants were routinely quantified by the BCA method (BCA Protein Assay, Pierce) and stored at -20 or -80°C. Samples were then prepared with the lysates and Laemmli 4x solution containing 5% β-mercaptoethanol and boiled for 10 min. Samples were loaded onto precast SDS-PAGE gels and subjected to electrophoresis for resolution of the proteins of interest at room temperature using Tris-Glycine-SDS running buffer in a constant electric field of 100 V cm-1. Subsequently, proteins were transferred to polyvinylidene difluoride (PVDF) membranes by transfer with Tris-Glycine-Ethanol buffer at constant 35 mA overnight. The membranes were blocked for 1 h at room temperature in the blocking buffer (TBS1X with 5 % milk or BSA and 0.1 % Tween-20). Then, the membranes were incubated with primary antibodies 1 or 2 h at RT in blocking buffer. After washing 3 times for 5 min with 0.5 % TBS-Tween 20, the membranes were incubated for 1 h at RT with HRP-conjugated secondary antibodies in blocking buffer. Detection was performed using ECL (GE Healthcare). Images were captured with Fusion (Vilber Lourmat) and quantified with ImageJ software (NIH).

### Inclusion Forming Unit Analysis (IFU)

Inclusion-forming unit (IFU) analysis was performed as described previously (64). Briefly, cells were incubated for 48 h or 72, as indicated, lysed by physical destruction with tip and stored in SPG buffer. Then, dilutions were inoculated in serial dilutions of cell lysate into HeLa cells seeded in 96-well plates. After 24-48 h of secondary infection, cells were fixed, permeabilized and stained with FITC-coupled anti-MOMP antibodies. In the case of using GFP-overexpressing bacteria, only their observation by microscopy was performed.

Inclusions were visualized and counted in 30 fields. Image acquisition was performed on an inverted wide-field DMI6000 microscope (Leica Microsystems, Wetzlar) with a 100x (1.4 NA) objective and an Orca Flash 4LT+ camera (Hamamatsu Photonics). Z series of images were taken at 0.3 μm z-step increments. Images were also acquired with an ImageXpress Micro (Molecular Devices, Sunnyvale, USA).

In the case of IFU analyses relativized to INPUT, prior quantification of the number of inclusions that gave rise to inclusion-forming units was performed. That is, primary infection was performed in duplicate, one replicate was fixed and used to quantify inclusions (INPUT), while the other was lysed and the infective progeny collected for subsequent quantification (OUTPUT).

### Immunofluorescence

Formaldehyde-fixed cells were incubated with 50 mM NH Cl in PBS1X, and permeabilized in 2% FCS and 0.05% saponin PBS (permeabilization buffer). The cells were then incubated with the primary antibody for 1 h in the same buffer, washed in the permeabilization buffer and incubated with secondary antibodies coupled to FITC, Cy3 or Cy5 (Jackson ImmunoResearch and Invitrogen) in the same buffer for 45 min. Cells were then washed 3 times in permeabilization buffer, incubated in DiAmidino Phenyl Indole (DAPI) for 5 min before mounting the coverslips in Fluoromount G (Thermo Fisher Scientific). Immunofluorescence images were acquired on an inverted wide-field DMI6000 microscope (Leica Microsystems, Wetzlar) with a 100x (1.4 NA) objective and an Orca Flash 4LT+ camera (Hamamatsu Photonics). Images were processed with Adobe Illustrator CS5 (Adobe Systems, Inc., San Jose, CA, United States) and MacBiophotonics ImageJ.

### HIV/CT co-infection in human macrophages

Human macrophages were co-infected with HIV and CT in two ways: i) acute infection and ii) established infection.

1. Acute infection: monocyte-derived macrophages obtained as mentioned above were initially infected with CT-GFP at an MOI of 1 as previously described (centrifugation, 30 min, 1000g, 10°C) late in the day. On the morning of the next day, infection was continued with HIV WT at an MOI of 1. For this, macrophages were incubated with HIV or Mock for 6 hours and then the cells were washed with PBS 3 times before the medium was renewed. Samples were then collected at 24, 48 and 72 hours after the onset of HIV infection for different techniques (WB, IF, ELISA, IFU, TZM-bl).
2. established infection: Macrophages were incubated with HIV or Mock at an MOI of 1 for 7 days. Subsequently, the pre-infected macrophages were superinfected with CT-GFP or Mock at an MOI of 1 for 24 or 48 h. Samples were collected for analysis by various techniques.

### ELISA CAp24

The determination of HIV CAp24 viral protein expression inside macrophages and in the macrophage culture supernatant was determined by ELISA (enzyme-linked immunosorbent assay) with a commercial kit Alliance HIV-1 p24 ANTIGEN ELISA Kit 96 wells (PerkinElmer). The samples used were supernatants, which were stored at -80°C, and cell lysates (obtained in the same manner as for WB and stored at -20°C). In cases where the infection was chronic, a 1/50 pre-dilution had to be performed, whereas in acute infection no dilution was performed. For the absolute quantification of p24, a calibration curve was performed according to the manufacturer’s instructions. The reaction was inactivated with 4N sulfuric acid and the color generated was measured according to the absorbance at 450 nm in an Multiskan FC plate spectrophotometer (Thermo Scientific).

### Cytotoxicity assay

The LDH (lactate dehydrogenase) enzyme activity in the supernatant of uninfected, infected and co-infected human macrophages was determined with the commercial Thermo Scientific™ Pierce™ LDH Cytotoxicity Assay Kit. Supernatants from cultured macrophages were taken at the indicated post-infection times and subsequently stored at - 80°C prior to processing. Determination was performed by end-point colorimetric reaction and then absorbance determination on Multiskan FC plate spectrophotometer (Thermo Scientific) at 505 nm as indicated in the manufacturer’s protocol. Because the experiment was not performed for the purpose of using this particular kit, the 100% cell lysis control, which is necessary for quantification in terms of percentage viability, was missing. It is for this reason that the results of this experiment are expressed in absorbance values and not as a percentage. The data were processed in GraphPad PRISM.

## ACKNOWLEDGMENTS

We thank Dr Agathe Subtil (Institut Pasteur Paris) for kindly providing the CT strains and Dr Cecile Arrieumerlou for her help with image acquisition on the Molecular Devices microscope. We thank the IMAG’IC facility of Institut Cochin that is part of the national France-BioImaging infrastructure supported by Agence Nationale de la Recherche (ANR-10-INBS-04). Work in the laboratory of F.N. was supported by grants from CNRS, INSERM, Université Paris Cité. Work in the laboratory of MTD was supported by PICT 2020 and SIIP-UNCUYO. Collaborative work in the laboratories of FN and MTD was supported by visiting programs supported by the Centro Franco Argentino de Cuyo, the French Embassy in Argentina, the Universidad Nacional de Cuyo, and Université Paris Cité. MAB was the recipient of PhD fellowship from CONICET and a “Saint Exupery” fellowship supported by the French “Ministère de l’Europe et des Affaires Etrangères” and the Argentinian “Ministerio de Educacion, Cultura, Ciencia y tecnologia”.

